# *Taenia* larvae possess distinct acetylcholinesterase profiles with implications for host cholinergic signalling

**DOI:** 10.1101/2020.06.12.148007

**Authors:** Anja de Lange, Ulrich Fabien Prodjinotho, Hayley Tomes, Jana Hagen, Brittany-Amber Jacobs, Katherine Smith, William Horsnell, Chummy Sikasunge, Murray E. Selkirk, Clarissa Prazeres da Costa, Joseph Valentino Raimondo

## Abstract

Larvae of the cestodes *Taenia solium* and *Taenia crassiceps* infect the central nervous system of humans. *Taenia solium* larvae in the brain cause neurocysticercosis, the leading cause of adult-acquired epilepsy worldwide. Relatively little is understood about how cestode-derived products modulate host neural and immune signalling. Acetylcholinesterases, a class of enzyme that degrade acetylcholine, are produced by a host of parasitic worms to aid their survival in the host. Acetylcholine is an important signalling molecule in both the human nervous and immune systems, with powerful modulatory effects on the excitability of cortical networks. Therefore, it is important to establish whether cestode derived acetylcholinesterases may alter host neuronal cholinergic signalling. Here we make use of multiple techniques to profile acetylcholinesterase activity in different extracts of both *Taenia crassiceps* and *Taenia solium* larvae. We find that the larvae of both species contain substantial acetylcholinesterase activity. However, acetylcholinesterase activity is lower in *Taenia solium* as compared to *Taenia crassiceps* larvae. Further, whilst we observed acetylcholinesterase activity in all fractions of *Taenia crassiceps* larvae, including on the membrane surface and in the excreted/secreted extracts, we could not identify acetylcholinesterases on the membrane surface or in the excreted/secreted extracts of *Taenia solium* larvae. Finally, using whole-cell patch clamp recordings in rat hippocampal brain slice cultures, we demonstrate that *Taenia* larval derived acetylcholinesterases can modify neuronal responses to acetylcholine. Together, these findings highlight the possibility that *Taenia* larval acetylcholinesterases can interfere with cholinergic signalling in the host, potentially contributing to pathogenesis in neurocysticercosis.

**Author summary:** Infection of the human nervous system with larvae of the parasite *Taenia solium* is a significant cause of acquired epilepsy worldwide. Despite this, the precise cellular and molecular mechanisms underlying epileptogenesis in neurocysticercosis remain unclear. Acetylcholinesterases are a family of enzymes widely produced by helminthic parasites. These enzymes facilitate the breakdown of acetylcholine, which is also a major neurotransmitter in the human nervous system. If *T. solium* larvae produce acetylcholinesterases, this could potentially disrupt host cholinergic signalling, which may in turn contribute to seizures and epilepsy. We therefore set out to investigate the presence and activity of acetylcholinesterases in *T. solium* larvae, as well as in *Taenia crassiceps* larvae, a species commonly used as a model parasite in neurocysticercosis research. We found that both *T. crassiceps* and *T. solium* larvae produce acetylcholinesterases with substantial activity. We further demonstrate that the acetylcholinesterase activity in the products of these parasites is sufficient to disrupt cholinergic signalling in an ex vivo brain slice model. This study provides evidence that *Taenia* larvae produce acetylcholinesterases and that these can interfere with cholinergic signalling in the host and potentially contribute to pathogenesis in neurocysticercosis.

## Introduction

Neurocysticercosis is a human disease which arises when larvae of the cestode *Taenia solium (T. Solium)* infect the central nervous system (1). The most common symptom of this infection is the development of epileptic seizures, which occur in 70–90% of symptomatic neurocysticercosis cases (2). As a result, neurocysticercosis is a major cause of adult-acquired epilepsy worldwide. Neurocysticercosis impacts heavily on the quality of life of those infected, and is also a significant drain on the medical and economic resources of endemic countries (3–5). Despite the global impact of neurocysticercosis, precisely how cerebral infection with *T. solium* relates to the development of seizures remains unclear.

It has been well documented that many parasitic worms of the alimentary tract produce substances that aid them in modulating host responses in ways that benefit the parasite (6–8). Acetylcholinesterases (AChEs), which catalyse the breakdown of acetylcholine, are one family of enzymes that have been implicated in the modulation of host responses. Helminths widely express membrane-bound forms of AChEs, which are classically associated with the facilitation of rapid acetylcholine signalling to parasite muscle, sensory, and neural structures (9,10). Some also produce surface-presenting membrane-bound AChEs (11–14), or can actively excrete/secrete AChEs, which may modulate acetylcholine dependent components of the host immune response, play a role in detoxification of ingested cholinesterase inhibitors, or inhibit smooth muscle contraction, mucus and fluid secretion associated with clearance of intestinal parasites (10,15–17).

Acetylcholine is also a major neurotransmitter in the human brain, with powerful effects on the excitability of cortical circuits (18,19). It is a critical component of multiple brain systems that are responsible for functions such as attention, learning, memory, sleep and motor activity (20,21). Disruption of cholinergic signalling is well known to lead to seizures. Amongst others, mutations of the nicotinic acetylcholine receptor underlies a heritable form of epilepsy (22), and pilocarpine (an acetylcholine muscarinic receptor agonist) is a well described proconvulsant agent (23). Further, blockade of endogenous brain AChEs by organophosphate pesticides or poisons can also lead to seizures (24,25).

Since *T. solium* larvae invade the central nervous system in neurocysticercosis, it is important to determine potential AChE activity expressed by these larvae, as such activity could conceivably interfere with endogenous cholinergic signalling in the brain. *Taenia crassiceps* (*T. crassiceps*) is a related cestode, which has also been known to invade the human nervous system, and is widely utilised as a model parasite for *T. solium* in neurocysticercosis research (26,27). It is therefore also important to ascertain how AChE activity might compare between the larvae of these two *Taenia* species.

AChEs have been reported in the adult forms of several members of the broader *Taeniidae* family (28–30) as well as in larval stages (9,11,13,31,32). The presence of AChEs in metacestodes of *Echinococcus granulosus* is particularly noteworthy, as these are known to infect the nervous system of children (33). The AChEs are often associated with the neural structures and parasite tegument of Taeniid*s*, and there is also some suggestion that some Taeniid larvae may release AChEs into the host environment (31). Studies describing cholinesterases in *T. crassiceps* larvae are scarce, with one report of AChEs in the bladder wall of T. crassiceps (34), and one other study which refers to the presence of “unidentified esterases” in the cystic fluid of *T. crassiceps* (35).

A histological study by Vasantha *et al.* (36) in *T. solium* larvae demonstrated AChE staining in neural structures of the larvae. No obvious staining of AChEs on the surface of the larvae is described, apart from positive staining in a few surface nerve endings. We further found one other report of cholinesterase activity in *T. solium* larvae, with activity predominantly present in the isolated cyst bladder ((37) cited in (38)).

Therefore, there is an important need for a detailed characterization of AChE activity in the larvae of different *Taenia* species, as well as an investigation into whether larval derived AChEs could conceivably disrupt host neuronal cholinergic signalling. Here, we used multiple techniques to explore AChEs activity in different extracts of both *T. crassiceps* and *T. solium* larvae. We find that both the larvae of *T. crassiceps* and *T. solium* contain significant AChE activity, but it is broadly lower in *T. solium* as compared to *T. crassiceps* larvae. In addition, whilst AChEs were present in all fractions of *T. crassiceps* larvae, including the membrane surface and excreted/secreted extracts, we could not identify AChEs on the membrane surface or within the excreted/secreted extracts of *T. solium* larvae. Finally, using whole-cell patch clamp recordings in rodent hippocampal brain slice cultures we demonstrate that *Taenia* larval derived AChEs can modify neuronal responses to acetylcholine.

## Materials and Methods

### Ethics statement

All animal handling, care and procedures were carried out in accordance with South African national guidelines (South African National Standard: The care and use of animals for scientific purposes, 2008) and with approval from the University of Cape Town Animal Ethics Committee (Protocol No: AEC 019/025, AEC 014/035).

### *Taenia* acquisition, maintenance, and preparation of cyst extracts

#### Acquisition and maintenance of T. crassiceps larvae

Larvae (ORF strain) were donated by Dr Siddhartha Mahanty (University of Melbourne, Melbourne, Australia) and propagated *in vivo* by serial intraperitoneal infection of 5-8-week-old female C57BL/6 mice. Every 3 months parasites were harvested by peritoneal lavage and washed 6 times in phosphate buffered saline (PBS, 1X, pH 7.4) before further processing.

#### Preparation of T. crassiceps whole cyst homogenate

Larvae were frozen at −80 °C immediately after harvesting. Upon thawing, larvae were suspended in a volume of PBS threefold that of the larvae. A protease inhibitor cocktail was added to this suspension (1% vol/vol, Sigma-Aldrich). The larvae were then homogenised on ice using a glass tissue grinder. The resulting mixture was centrifuged at 3100 *g* for 20 minutes at 4 °C. The liquid supernatant (excluding the low density white floating layer) was collected and sterile filtered through a 0.22 μm size filter (Millex-GV syringe filter, Merck). This supernatant was then aliquoted and stored at −80 °C until use. This preparation is referred to as *“T. crassiceps* whole cyst homogenate”.

#### Preparation of T. crassiceps cyst membrane and cyst vesicular fluid extracts

After harvesting, washed larvae (+/− 10ml) were placed onto a piece of filter paper (which had been saturated with 1X PBS) in a metal sieve. Cysts were then ruptured using a weighing spatula. The fluid from the ruptured cysts that passed through the filter paper was collected in a beaker and was centrifuged at 3100 *g* for 20 minutes at 4 °C, and the supernatant was collected, aliquoted and stored at −80 °C until use. This extract is referred to as *“T. crassiceps* cyst vesicular fluid”. The fraction of the cysts that remained on the filter paper were scraped off with the weighing spatula and suspended in an equal volume of PBS containing a protease inhibitor cocktail (1% vol/vol, Sigma-Aldrich). This mixture was freeze-thawed once at −80 °C, homogenised on ice using a glass tissue grinder, and centrifuged at 3100 *g* for 20 minutes at 4 °C. The liquid supernatant was collected, aliquoted, and stored at −80 °C until use. This extract is referred to as *“T. crassiceps* cyst membrane”.

#### Preparation of T. crassiceps larval excretory/secretory extracts

After harvesting, washed larvae (+/− 10 ml) were placed in a 50 ml culture flask with 10 ml culture medium (Earle’s Balanced Salt Solution with 5.3 g/L glucose, 1X Glutamax, 50 U/ml penicillin, 50 μg/ml streptomycin, 100 μg/ml gentamicin sulphate and 11.4 U/ml nystatin). Larvae were maintained at 37 °C in 5 % CO_2_. After 48 hrs the medium was discarded and replaced with 10 ml fresh media. At 20 days *in vitro* (at which point larvae still displayed motility) the culture media was collected, aliquoted, and stored at −80 °C until use. For electrophysiology experiments the excretory/secretory extracts were dialysed/buffer exchanged to PBS using an Amicon stirred cell (Merck) with a 3 kDa molecular weight cut-off membrane, in order to remove small molecules that could potentially induce electrophysiological responses that would interfere with the acetylcholine effect (such as glutamate – see Tomes *et al.,* 2020).

#### Acquisition of T. solium larvae

Larvae of *T. solium* were harvested from the muscles of a heavily infected, freshly slaughtered pig in Lusaka, Zambia. Larvae were removed from the muscle by vigorous shaking and collected in petri dishes containing sterile PBS (1X, pH 7.4).

#### Preparation of T. solium whole cyst homogenate

After extensive washing with sterile PBS (1X, pH 7.4), larvae were suspended in a volume of PBS threefold that of the larvae, containing phenylmethyl-sulphonyl fluoride (5 mM) and leupeptin (2.5 μM). Larvae were then homogenised using a sterile handheld homogenizer at 4 °C. The resulting homogenate was sonicated (4 x 60 s at 20 kHz, 1 mA, with 30 s intervals), gently stirred with a magnetic stirrer (2 hrs at 4°C), and centrifuged at 15 000 *g* for 60 min at 4 °C. The liquid supernatant (excluding the low density white floating layer) was collected and sterile filtered through 0.45 μm size filters (Millex-GV syringe filter, Merck). This supernatant was then collected, aliquoted and stored at −80 °C until use. This preparation is referred to as *“T. solium* whole cyst homogenate”.

#### Preparation of T. solium cyst vesicular fluid extracts and cyst membrane and scolex

After extensive washing with sterile PBS, larvae were placed in a petri dish and individually ruptured with a sterile needle. The resulting fluid in the petri dish was collected and centrifuged at 15 000 *g* for 60 min at 4 °C. The supernatant was then sonicated (4 x 60 s at 20 kHz, 1 mA, with 30 s intervals), phenylmethyl-sulphonyl fluoride (5 mM) and leupeptin (2.5 μM) were added, and the solution was centrifuged a second time at 15,000 *g* for 60 min at 4 °C. The supernatant was collected, aliquoted and stored at −80 °C until use. This extract is referred to as *“T. solium* cyst vesicular fluid”. The remaining parts of the larvae were again extensively washed with PBS and then suspended in an equal volume of PBS containing phenylmethyl-sulphonyl fluoride (5 mM) and leupeptin (2.5 μM). This suspension was again homogenised using a sterile handheld homogenizer at 4 °C. The resulting homogenate was sonicated (4 x 60 s at 20 kHz, 1 mA, with 30 s intervals), gently stirred with a magnetic stirrer (2h at 4°C), and centrifuged at 15,000 *g* for 60 min at 4 °C. The liquid supernatant (excluding the low density white floating layer) was collected and sterile filtered through 0.45 μm size filters (Millex-GV syringe filter, Merck). This supernatant was then aliquoted and stored at −80 °C until use. This extract is referred to as “*T. solium* cyst membrane and scolex”.

#### Preparation of T. solium excretory/secretory extracts

After harvesting, washed larvae were placed into 6 well plates (+/− 15 per well) with 2 ml culture medium (RPMI 1640 with 10 mM HEPES buffer, 100 U/ml penicillin, 100 μg/ml streptomycin, 0.25 μg/ml amphotericin B and 2 mM L-glutamine). Every 24 h, 1 ml of culture medium was collected from each well and replaced with fresh culture medium. Medium from all wells was pooled each day, aliquoted and stored at −80 °C. Media collected on days 1, 2, and 3 *in vitro* were pooled, and are referred to as *“T. solium* excretory/secretory extracts”.

All *T. crassiceps* and *T. solium* larval extracts were assessed for protein concentration using a BCA or Bradford protein assay kit (Sigma-Aldrich), respectively.

### Acetylcholinesterase activity and inhibitor sensitivity

AChE activity was determined by the method of Ellman *et al.* (40) at room temperature with 1 mM acetylthiocholine iodide as substrate in the presence of 1 mM 5,5’-dithiobis(2-nitrobenzoic acid) (DTNB) in 100 mM sodium phosphate (pH 7.0). The reaction was monitored by measuring the absorbance at 412 nm, and hydrolysis of acetylthiocholine iodide calculated from the extinction coefficient of DTNB (40). Activity was expressed as nanomoles of acetylthiocholine hydrolysed per minute per milligram of total protein in each larval extract (nmol min^-1^ mg^-1^). To test the sensitivity of *Taenia* AChEs to different inhibitors, extracts were preincubated with different concentrations of 1,5-bis(4-allyldimethylammoniumphenyl)pentan-3-one dibromide (BW 284c51), tetraisopropyl pyrophosphoramide (iso-OMPA) or eserine salicylate for 20 min at room temperature in Ellman buffer, prior to the addition of 1 mM acetylthiocholine iodide and enzyme activity determination. Each reaction was assayed a minimum of three times. Where AChE activity was reduced to undetectable levels by inhibitors, a residual activity of 0 % was allocated on inhibition curves.

### Non-denaturing polyacrylamide gel electrophoresis (PAGE)

Extracts were electrophoresed in Tris-glycine buffer, pH 8.3, through 7.5 % polyacrylamide gels in the absence of denaturing and reducing agents. Electrophoresis was performed at 150 V for 3 hrs on ice. Protein staining (Coomassie) was performed on one set of PAGE gels, and specific staining for AChE activity was performed overnight as described by Selkirk and Hussein (41) adapted from the method of Karnovsky and Roots (42). The maximum volume of each protein extract was loaded (20 μl), to ensure maximal staining. Protein concentrations of the different extracts varied (*T. crassiceps:* whole cyst homogenate = 1.9 mg/ml, cyst membrane = 3.4 mg/ml, cyst vesicular fluid = 3.0 mg/ml and excretory/secretory extracts = 1.32 mg/ml; *T. Solium:* all extracts = 1.5 mg/ml). Each stain was performed at least three times to confirm reproducibility.

### *In situ* localisation of acetylcholinesterases

To localise *Taenia* AChEs, fresh *Taenia* larvae were submerged in 10 % formalin for 60 min, to fix the tissue. Some of the larvae were then stained overnight for AChE activity as described by Selkirk and Hussein (41) adapted from the method of Karnovsky and Roots (42), and mounted onto slides as whole mounts. A subset of the fixed larvae was embedded in cryo embedding medium, frozen overnight at −80 °C, and cryo-sectioned the following day at 50 μm. The sections were then similarly stained overnight for AChE activity, placed on positively charged slides and dehydrated in graded alcohols before mounting. To assess non-specific staining, in a subset of the whole mount and cryo-section specimens, acetylthiocholine iodide (the substrate) was omitted during the AChE staining procedure. Specimens were imaged using an upright light microscope.

### An *ex vivo* model to examine the effect of *Taenia* AChEs in the context of neurocysticercosis

#### Hippocampal brain slice preparation

Organotypic brain slices were prepared using 6-8-day-old Wistar rats following the protocol originally described by Stoppini *et al.* (43). Briefly, brains were extracted and swiftly placed in cold (4°C) dissection media consisting of Earle’s Balanced Salt Solution (Sigma-Aldrich) supplemented with D-glucose (6.1 g/L) and HEPES (6.6 g/L). The hemispheres were separated, and individual hippocampi were removed and immediately cut into 350 μm slices using a Mcllwain tissue chopper (Mickle). Cold dissection media was used to separate and rinse the slices before placing them onto Millicell-CM membranes (Sigma-Aldrich). Slices were maintained in culture medium consisting of 25 % (vol/vol) Earle’s balanced salt solution; 49 % (vol/vol) minimum essential medium (Sigma-Aldrich); 25 % (vol/vol) heat-inactivated horse serum (Sigma-Aldrich); 1 % (vol/vol) B27 (Invitrogen, Life Technologies) and 6.2 g/l D-glucose (Sigma-Aldrich). Slices were incubated in a 5 % carbon dioxide (CO_2_), humidified incubator at 37 °C. Recordings were made after 6-14 days in culture.

#### Electrophysiology

Brain slices were transferred to a submerged recording chamber on a patch clamp rig, which was maintained at a temperature between 28 and 34 °C, and were continuously superfused with standard artificial cerebrospinal fluid (120 mM NaCl, 3mM KCl, 2 mM MgCl_2_, 2 mM CaCl_2_, 1.2 mM NaH_2_PO_4_, 23 mM NaHCO_3_ and 11 mM D-Glucose in deionised water with pH adjusted to between 7.35 – 7.40 using 0.1 mM NaOH) bubbled with carbogen gas (95 % O2: 5 % CO_2_) using peristaltic pumps (Watson-Marlow). Micropipettes were prepared (tip resistance between 3 and 7 MΩ) from borosilicate glass capillaries (outer diameter 1.2 mm, inner diameter 0.69 mm) (Harvard Apparatus Ltd) using a horizontal puller (Sutter). Micropipettes utilised for whole cell patch clamping were filled with an artificial cell internal solution (126 mM K-gluconate, 4 mM KCl, 10 mM HEPES, 4 mM Na2ATP, 0.3 mM NaGTP and 10 mM Na2-phosphocreatine) before being placed over the recording electrode.

Neurons in the CA3 region of the hippocampus were visualized using an upright microscope with a 20X water immersion objective. Surface cells with a typical pyramidal cell body morphology were selected for whole cell patching. To demonstrate the effect of acetylcholine on hippocampal CA3 pyramidal neurons a second micropipette containing a solution of 200 μm acetylcholine was lowered to the cell surface once a cell had been patched. Current was injected to hold the membrane potential of cells close to their action potential firing threshold and then five 30 ms puffs (~20 psi) of the solution was applied to the cell’s surface using an OpenSpritzer (Forman *et al.,* 2017) and the neuron’s response recorded for 26 s before a 94 s “recovery” period was allowed. This cycle was performed 5 times.

To explore the ability of *Taenia* acetylcholinesterase activity to alter neuronal acetylcholine signalling, another set of patch-clamp experiments were performed, this time with two “puffer” micropipettes being lowered to the cell surface once a neuron had been patched. One of these micropipettes contained a solution of 200 μm acetylcholine with 1.3 mg/ml *T. crassiceps* excretory/secretory extracts, while the second contained a solution of 200 μm acetylcholine with 1.3 mg/ml *T. crassiceps* excretory/secretory extracts that had been heated to 56 °C for 30 min to inactivate enzymes. Again, current was injected to hold the membrane potential of cells close to their action potential firing threshold. Five 30 ms puffs (~20 psi) of one of the solutions was then applied to the cell’s surface and the neuron’s response recorded for 26 s before a 94 s “recovery” period was allowed. Thereafter an identical puff train of the other solution was applied, and the neuron’s response again recorded for 26 s. After another 94 s recovery period, the cycle was repeated.

Post-recording analysis consisted of counting the number of action potentials induced by the application of each solution within a 5 s period of the onset of the puff train. Only traces with a baseline membrane potential prior to the puff application of between −60 mV and −45 mV were included. Each data point in the puffing experiments represents the average of between 2 and 5 repeats of the puff cycle. Matlab (MathWorks) was utilised for trace analysis.

### Data analysis and statistics

Data was visualised and analysed using Matlab, Microsoft Excel and GraphPad Prism. Each dataset was subjected to a Shapiro-Wilk test to determine whether it was normally distributed. Most datasets proved to be non-normal and as such non-parametric analyses were utilised throughout. These included: Kruskal-Wallis analyses with Dunn’s multiple comparison post-hoc tests and Mann-Whitney tests. The confidence interval for all tests was set at 95 %.

## Results

### All *T. crassiceps* larval extracts display acetylcholinesterase activity

In order to quantify AChE activity in *T. crassiceps* larvae, Ellman’s assays were employed, using acetylthiocholine as a substrate (40). These assays revealed that all *T. crassiceps* larval extracts had significant AChE activity (**Table 1, Fig. 1A**). A Kruskal Wallis one-way ANOVA with post hoc Dunn’s Multiple Comparison tests revealed that the only statistically significant difference between the median activities of the different *T. crassiceps* larval extracts was between that of the cyst vesicular fluid and that of the excretory/secretory extracts (P ≤ 0.01, **Fig. 1A**).

**Table 1:** Acetylcholinesterase (AChE) activity of different larval extracts of T. crassiceps and T. solium

| Larval species | Larval extract | # of assays | Median AChE activity (nmol min<sup>-1</sup> mg<sup>-1</sup>) |
| --- | --- | --- | --- |
| ***Taenia crassiceps*** | Whole cyst homogenate | 10 | 31.7 (IQR 24.0 – 58.3) |
|  | Cyst membrane | 10 | 39.7 (IQR 36.1 – 46.3) |
|  | Cyst vesicular fluid | 10 | 51.5 (IQR 50.9 – 54.1) |
|  | Excretory/secretory extracts | 10 | 31.8 (IQR 50.9 – 54.1) |
| ***Taenia Solium*** | Whole cyst homogenate | 5 | 4.1 (IQR 4.1 – 5.1) |
|  | Cyst membrane & scolex | 5 | 14.0 (IQR 13.8 – 15.3) |
|  | Cyst vesicular fluid | 5 | 4.7 (IQR 3.37 – 5.03) |
|  | Excretory/secretory extracts | 4 | Undetectable |

**Figure 1:**
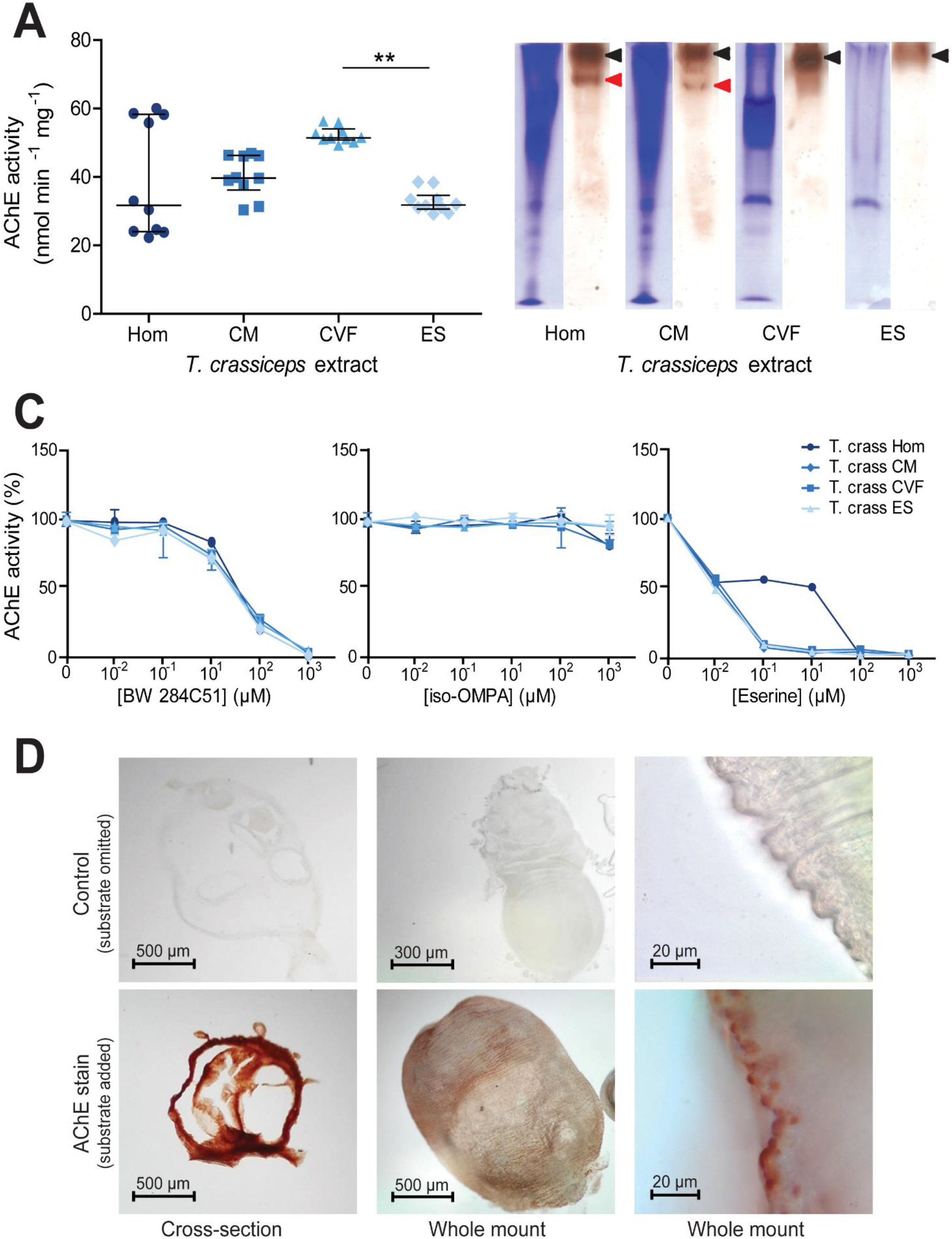
Identification and characterisation of acetylcholinesterases in *Taenia crassiceps* larval extracts. A) Quantification of acetylcholinesterase (AChE) activity in different *Taenia crassiceps* larval extracts. AChE activity was quantified using the method of Ellman *et al.* (40) with 1 mM acetylthiocholine iodide as substrate in the presence of 1 mM 5,5’-dithiobis(2-nitrobenzoic acid) in 100 mM sodium phosphate, pH 7.0, at room temperature. The extracts assessed were: whole cyst homogenate (Hom); cyst membrane (CM), cyst vesicular fluid (CVF) and larval excretory/secretory extracts (ES). Values with median ± IQR, N = 10 for all extracts assayed, **p ≤ 0.01, Kruskal-Wallis test with Dunn’s multiple comparison post-hoc tests B) Non-denaturing polyacrylamide gel electrophoresis of *Taenia crassiceps* extracts. Extracts were electrophoresed in Tris-glycine buffer, pH 8.3, through 7.5% polyacrylamide gels in the absence of denaturing and reducing agents. Coomassie staining was performed on one set of gels (left tracks), and staining for AChE activity (42) was performed on another set of gels for 16 hrs after incubation with the substrate (right tracks). The maximum volume of each extract was loaded (20 μl), to ensure maximal staining. Protein concentrations of the different extracts varied: Hom = 1.9 mg ml^-1^, CM = 3.4 mg ml^-1^, CVF = 3.0 mg ml^-1^ and ES = 1.32 mg ml^-1^. C) Inhibitor sensitivity of *Taenia crassiceps* AChEs. *Taenia crassiceps* extracts were preincubated with BW 284C51, iso-OMPA or eserine salicylate for 20 min at room temperature in Ellman buffer, prior to the addition of 1 mM acetylthiocholine iodide and enzyme activity determination. Median ± Range, N = 10 for all extracts in absence of inhibitors, N = 3 for all extracts at all inhibitor concentrations. D) Localisation of larval AChEs. Cryo-sections and whole mounts of *Taenia crassiceps* larvae were subjected to AChE staining (42) for 16 hrs prior to dehydration and mounting. Images on the left show time-matched controls where acetylthiocholine iodide was omitted from the staining solution.

To visually confirm *T. crassiceps* AChE activity, and to assess whether there may be more than one AChE isoform in *T. crassiceps* larval extracts, non-denaturing PAGE gels were run, and stained for AChE. A second set of non-denaturing PAGE gels were run simultaneously and Coomassie stained. These demonstrated that the different *T. crassiceps* larval extracts contained different protein compositions (left-hand tracks in **Fig. 1B**). The AChE stained gels (right-hand track for each larval extract in **Fig. 1B**) showed distinct dark bands (indicated by black arrowheads) in the tracks of all the *T. crassiceps* larval extracts, thereby confirming that all the *T. crassiceps* larval extracts show AChE activity. In the whole cyst homogenate and the cyst membrane tracks of the AChE stained gels, there was an additional smaller band (indicated by the red arrowheads in **Fig. 1B**). These results suggest that *T. crassiceps* larvae express more than one isoform of AChE.

### Inhibitor sensitivity of *T. crassiceps* larval acetylcholinesterases

The sensitivity of AChE activity in *T. crassiceps* larval extracts to different inhibitors was tested by preincubating them for 20 min with different concentrations of BW 284c51 (a selective AChE inhibitor), iso-OMPA (a selective butyryl cholinesterase inhibitor) or eserine salicylate (a nonselective cholinesterase inhibitor) before assaying AChE activity (40). All *T. crassiceps* extracts showed a similar dose-dependent inhibitory response to BW 284c51, with AChE activity in all extracts being almost completely inhibited by the presence of 1000 μM BW 284c51 (**Fig. 1C, S1 Table**). Conversely, the AChE activity of *T. crassiceps* extracts was not greatly inhibited by iso-OMPA, with only very small reductions in activity being observed even at high (1 mM) inhibitor concentration (**Fig. 1C, S1 Table**). AChE activity in *T. crassiceps* cyst membrane, cyst vesicular fluid and excretory/secretory extracts was highly sensitive to eserine inhibition, with strong inhibition apparent at low (1 μM) eserine concentration (**Fig. 1C, S1 Table**). The AChE activity of *T. crassiceps* whole cyst homogenate was less sensitive to eserine inhibition, only displaying strong inhibition at a much higher eserine concentration (100 μM) (**Fig. 1C, S1 Table)**. These inhibition patterns suggest that the cholinesterase produced by *T. crassiceps* larvae can be classified as true AChEs, as opposed to pseudocholinesterases (45).

### *T. crassiceps* acetylcholinesterases are ubiquitous within the tegument membrane and are present on the larval surface

To spatially localise AChEs within the larvae, both cross-sections of larvae and whole larvae were subjected to AChE staining (42), (**Fig. 1D**). To evaluate non-specific staining, a second set of larval cross-sections and whole larvae were subjected to the same staining procedure, with the exception that the substrate (acetylthiocholine iodide) was omitted. Samples where the substrate was omitted (**top row, Fig. 1D**) showed minimal staining. In contrast, cross sections stained for AChE activity displayed dense, uniform staining, indicating that AChEs are localised ubiquitously throughout the tegument membrane (**bottom-left panel, Fig. 1D**). Whole larvae stained for AChE showed light surface staining at low magnification (**bottom-centre panel, Fig, 1D**), and high magnification revealed that staining localised to numerous small protrusions on the surface of the cyst tegument membrane (**bottom-right panel, Fig, 1D**).

### *T. solium* larvae produce acetylcholinesterases but do not actively excrete/secrete them

Next, we focussed on the major pathogenic cestode of humans; *T. solium.* Ellman’s assays revealed that *T. solium* whole cyst homogenate, cyst membrane and scolex, and cyst vesicular fluid had detectable AChE activity, whilst the excretory/secretory extracts consistently displayed no detectable AChE activity (**Table 1, Fig. 2A**). A Kruskal Wallis one-way ANOVA with post hoc Dunn’s Multiple Comparison tests revealed that the cyst membrane and scolex had a statistically significantly higher activity than that of the whole cyst homogenate and the cyst vesicular fluid, whilst the median activity of two latter extracts did not differ significantly from one another (P ≤ 0.01, **Fig. 2A**).

**Figure 2:**
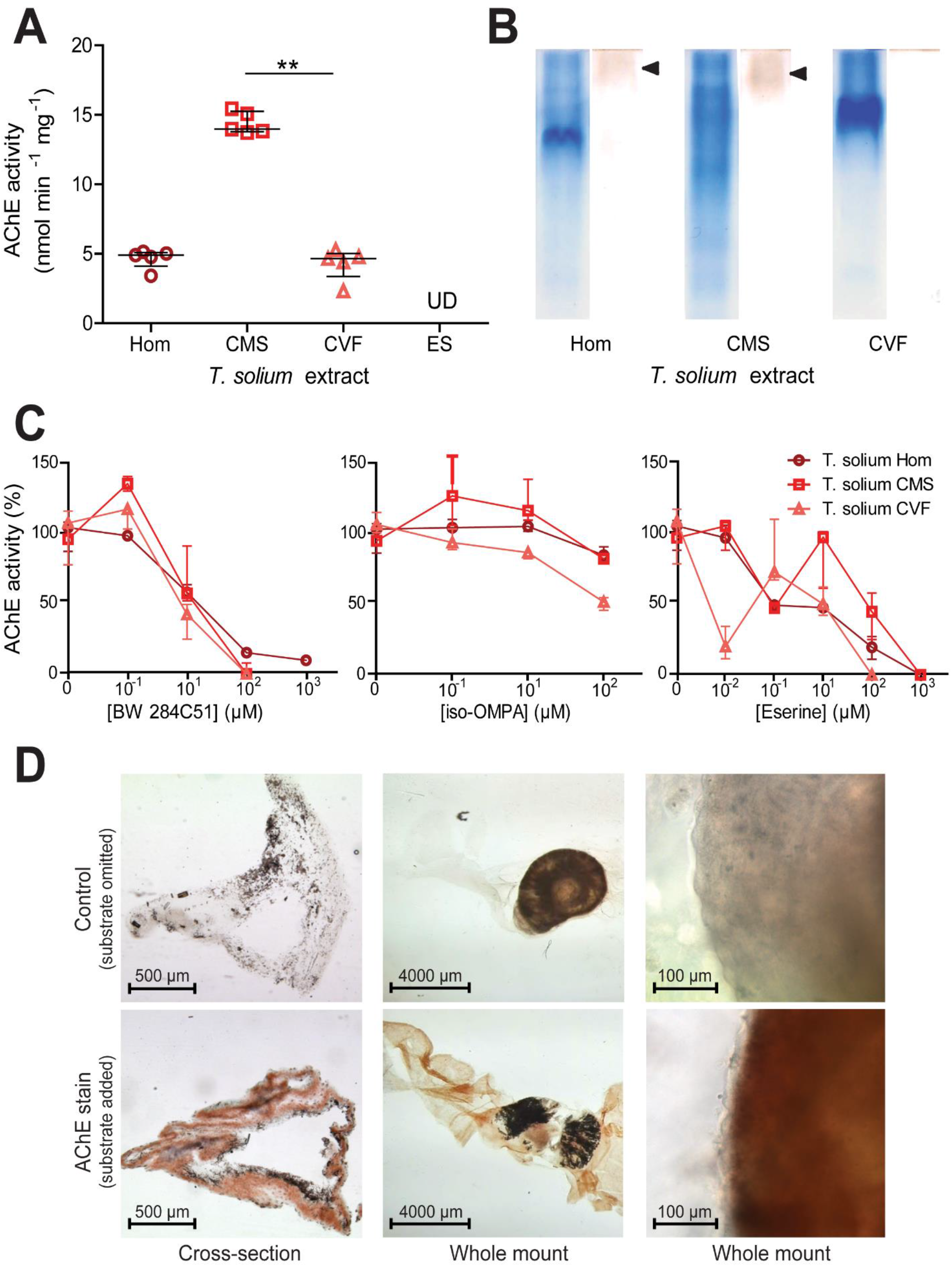
Identification and characterisation of acetylcholinesterases in *Taenia solium* extracts. A) Quantification of acetylcholinesterase (AChE) activity in different *Taenia solium* extracts as in Fig. 1. The extracts assessed were: whole cyst homogenate (Hom); cyst membrane and scolex (CMS), cyst vesicular fluid (CVF) and larval excretory/secretory extracts (ES). Values with median ± IQR, N = 5 for all extracts assayed, **p ≤ 0.01, Kruskal-Wallis test with Dunn’s multiple comparison post-hoc tests B) Non-denaturing polyacrylamide gel electrophoresis of different *Taenia solium* extracts. Coomassie staining was performed on one set of PAGE gels (left tracks), and staining for AChE activity (42) was performed on another set of gels (right tracks). For each extract, 30 μg of total protein was loaded. C) Inhibitor sensitivity of *Taenia solium* AChEs. *Taenia solium* extracts were preincubated with BW 284C51, iso-OMPA or eserine for 20 min at room temperature in Ellman buffer, prior to the addition of 1 mM acetylthiocholine iodide and enzyme activity determination. Median ± Range, N = 5 for all extracts in absence of inhibitors and at 10 μM inhibitor concentration, N = 3 for all extracts at all other inhibitor concentrations. D) Localisation of larval AChEs. Cryosections and whole mounts of *Taenia solium* larvae were subjected to AChE staining (42). Images on the left show time-matched controls where acetylthiocholine iodide was omitted from the staining solution.

To visually confirm AChE activity in *T. solium,* and to assess whether there may be more than one AChE isoform in larval extracts, non-denaturing PAGE gels were resolved and stained for AChE (Fig. 2B). Again, a second set of non-denaturing PAGE gels were run simultaneously and Coomassie stained. The Coomassie stain showed that each extract has a distinct protein profile composition (left-hand tracks in Fig. 2B). The AChE stained gels (right-hand track for each larval extract in Fig. 1B) revealed bands in the whole cyst homogenate and in the cyst membrane and scolex preparations (indicated by black arrowheads), but no apparent band in the cyst vesicular fluid track, likely due to insufficient enzyme concentration in the amount of extract resolved.

### Inhibitor sensitivity of *T. solium* larval acetylcholinesterases

The sensitivity of AChE activity in *T. solium* larval extracts to different inhibitors was tested by preincubating them for 20 min with different concentrations of BW 284c51, iso-OMPA or eserine salicylate, before assaying AChE activity (40). All *T. solium* extracts showed a similar dose-dependent inhibitory response to BW 284c51, although the *T. solium* whole cyst homogenate appears somewhat less sensitive to inhibition that the cyst membrane and scolex and cyst vesicular fluid (**Fig. 2C, S2 Table**). *T. solium* extracts showed low sensitivity to inhibition by iso-OMPA (**Fig. 2C, S2 Table**). *T. solium* whole cyst homogenate, cyst membrane and scolex, and cyst vesicular fluid showed very variable sensitivities to inhibition by increasing concentrations of eserine, but were ultimately all strongly inhibited at an eserine concentration of 1 mM or less (**Fig. 2C, S2 Table**). These inhibition patterns suggest that the cholinesterases produced by *T. solium* larvae can be classified as true AChEs, as opposed to pseudocholinesterases, although the fact that some inhibition is displayed at high iso-OMPA concentrations may suggest a small pseudocholinesterase component (45).

### *T. solium* larval acetylcholinesterases are localised within the cyst tegument membrane, but do not appear to present on the surface of the parasite

Next, we set out to spatially localise AChEs within the larvae. To do so both cross-sections of larvae and whole larvae were subjected to the same AChE staining as was applied to *T. crassiceps* larvae. Control samples where the substrate was omitted to evaluate non-specific staining (**top row, Fig. 2D**), showed some patchy black background staining. AChE stained cross-sections and whole mounts showed uniform, although not very dense, AChE staining throughout the tegument membrane (in addition to the black background staining) (**bottom-left and bottom-centre panel, Fig. 2D**). High magnification images of the surface of the tegument membrane in whole-mounted AChE-stained larvae revealed that, unlike in *T. crassiceps,* AChEs in the tegument membrane of *T. solium* are not surface-presenting (**bottom-right panel, Fig, 2D**).

### Larvae of *T. solium* have less acetylcholinesterase activity as compared to *T. crassiceps,* and *T. crassiceps* larvae excrete/secrete acetylcholinesterases, whilst *T. solium* larvae do not

Comparison of AChE activity in the extracts of *T. crassiceps* versus the comparable *T. solium* extracts show that *T. solium* extracts consistently have significantly lower AChE activities than those of *T. crassiceps* (P ≤ 0.001, Mann Whitney tests, **Table 1**). Further, a different pattern of AChE distribution within the cyst is observed between the two species – *T. solium* AChEs appear to be predominantly located in the cyst membrane and scolex, whilst *T. crassiceps* AChEs appear abundant in both the cyst membrane and in the cyst vesicular fluid and are additionally excreted/secreted (**Table 1**). *T. crassiceps* also displays surface presenting AChEs, whilst *T. solium* do not, as revealed by AChE staining of whole larvae (**Fig. 1D & Fig. 2D**). It is also noteworthy that *T. crassiceps* and *T. solium* larval extracts display different inhibitor sensitivities to BW 284c51, iso-OMPA and particularly to eserine (**Fig. 1C** versus **Fig. 2C**). This suggests that the two parasites produce different forms of AChE, although further investigation would be required to confirm this.

### *Taenia* larval acetylcholinesterases have sufficient activity to modify neuronal acetylcholine signalling *ex vivo*

In order to investigate what implications the presence of *Taenia* larval AChEs may have in the context of neurocysticercosis, 200 μM acetylcholine (known to induce depolarisation in hippocampal pyramidal neurons (46) was applied to neurons in hippocampal organotypic cultures, together with either heat-inactivated *T. crassiceps* excretory/secretory products, or active *T. crassiceps* excretory/secretory products. The response of the membrane potential of the neurons was measured using whole-cell patch-clamp recordings (see Materials and Methods, and **Fig. 3A, B**). When neurons were held at a voltage close to their action potential threshold and picolitre volumes of 200 μM acetylcholine with heat-inactivated excretory/secretory products were puffed toward the soma of the neurons, they depolarised and fired action potentials (APs) (median = 3.6 APs, IQR = 1.0 – 11.2 APs, N = 16, **Fig. 3B,C**). However, when 200 μM acetylcholine with active excretory/secretory products were applied to the same neurons just 2 min prior to/after this, the neurons did not show the same response, often firing no action potentials despite still being held at a voltage close to their action potential threshold (median = 0.2 APs, IQR = 0.0 – 1.7 APs, N = 16, P = 0.0017, Wilcoxon signed-rank test, **Fig. 3C**). This demonstrates that the AChEs in *T. crassiceps* larval excretory/secretory products have sufficient activity to modify acetylcholine signalling in brain tissue. This would hold true for *T. solium* cyst membrane and scolex AChEs, which break down acetylcholine at roughly half the rate of *T. crassiceps* larval excretory/secretory extracts (**Table 1**).

**Figure 3:**
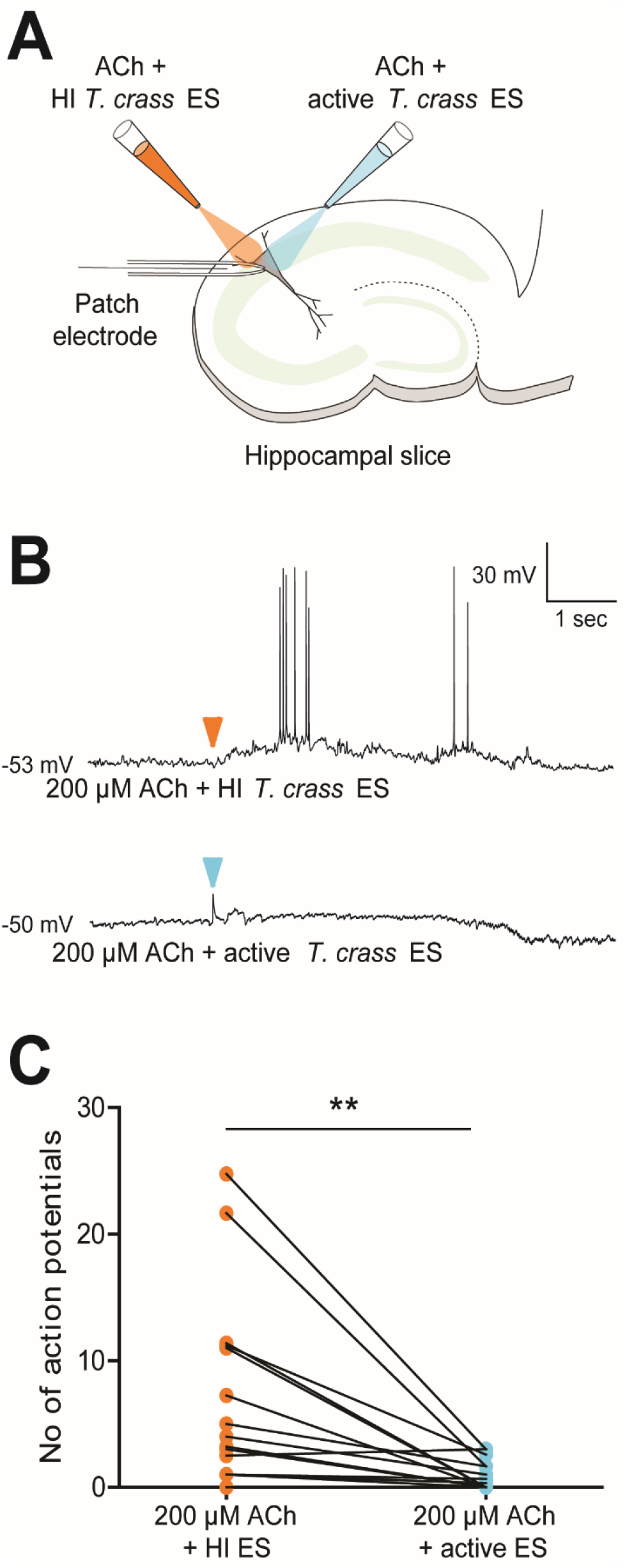
The functional effect of *Taenia* acetylcholinesterases on neuronal acetylcholine signalling. A) Schematic depicting the experimental setup where whole-cell patch-clamp recordings were made from rat CA3 pyramidal neurons in organotypic hippocampal brain slice cultures. Whilst recording the electrical activity from the neurons, two glass pipettes delivered picolitre volumes of either 200 μM acetylcholine with 1.3 mg ml^-1^ heat-inactivated *Taenia crassiceps* excretory/secretory extracts (orange pipette), or 200 μM acetylcholine with 1.3 mg/ml of standard *Taenia crassiceps* excretory/secretory extracts (blue pipette). B) The membrane potential responses of a pyramidal neuron when a solution of 200 μM acetylcholine with 1.3 mg ml^-1^ *Taenia crassiceps* excretory/secretory extracts that had been heat-inactivated at 56°C for 30 min (top trace) or 200 μM acetylcholine with 1.3 mg ml^-1^ unheated/active *Taenia crassiceps* excretory/secretory extracts (bottom trace) was puffed onto the cell body (arrowheads indicate moment of application). C) Population data (median ± IQR) where each point represents the mean number of action potentials evoked in 5 s after neurons were exposed to 5 x 30 ms puffs (2 – 5 cycles) of either a solution of 200 μM acetylcholine with 1.3 mg ml^-1^ heat-inactivated *Taenia crassiceps* excretory/secretory extracts (N = 13) or a solution of 200 μM acetylcholine with 1.3 mg ml^-1^ active *Taenia crassiceps* excretory/secretory extracts (N = 13). **p ≤ 0.01, Wilcoxon signed-rank test.

## Discussion

Here we have used multiple methods to characterise the amount and spatial localization of AChE activity in larvae of the cestodes *T. crassiceps* and *T. solium.* Previous studies have identified AChE activities in the larvae of multiple species of the broader Taeniidae family including *Echinococcus granulosus* (dog tapeworm) and *Taenia pisiformis* (rabbit tapeworm). To our knowledge our data represent the first definitive measurements of AChE activity from larvae of *T. crassiceps.* The amount of AChE activity we report in *T. crassiceps* whole cyst homogenate is similar to that previously reported for the larval homogenate of *T. pisiformis (T. crassiceps* whole cyst homogenate median activity = 31.69 nmol min^-1^ mg^-1^, T. pisiformis homogenate mean activity = 24.8 nmol min^-1^ mg^-1^)(13). Interestingly we found that whilst *T. solium* larvae also exhibit substantial AChE activity, this activity is broadly less than in extracts from *T. crassiceps.* Furthermore, the spatial profile of larval AChE activity is different between these two species. Whilst AChEs were present in all fractions of *T. crassiceps* larvae, and presented on the tegument membrane surface, we could not identify AChEs on the tegument membrane surface or within the excreted/secreted extracts of *T. solium* larvae. The lack of surface staining in *T. solium* larvae is in accordance with a previous study by Vasantha *et al.* (1992), which reports AChEs in *T. solium* larvae to be associated with a sub-tegumental network of nerves in the strobila and bladder wall.

Our findings have important implications in the context of *T. crassiceps* being utilised as a model organism for neurocysticercosis research, with both larval extracts and whole early-stage cysts of *T. crassiceps* being popular (47–50). The fact that *T. crassiceps* larvae have substantially higher AChE activity than *T. solium* larvae means that their potential to alter brain acetylcholinergic signalling is likely greater. The same would be true when utilising the tetrathyridia of *Mesocestoides corti* (another popular model parasite for neurocysticercosis research) as the reported AChE activity of these larvae is far greater than that which we report for both *T. crassiceps* and *T. solium* (51). The fact that *T. crassiceps* larvae express surface AChEs and actively excrete/secrete AChEs also has implications for research where whole, viable *T. crassiceps* larvae are utilised in neurocysticercosis models, as these *T. crassiceps* larvae may induce electrophysiological and immunological changes via AChE extraction that *T. solium* larvae may not.

Our observation that *T. solium* larvae do not excrete/secrete AChEs is interesting, as many helminth species have been observed to secrete these enzymes in substantial amounts, with proposed benefits to parasite survival, such as protection against ingested AChE inhibitors and modulation of the host immune response (7,10,16). Recently, a study by Vaux *et al.* (2016) demonstrated that *in vivo* exposure to secreted AChE from *Nippostrongylus brasiliensis* promoted classical activation of macrophages (as opposed to alternative activation), a state which is permissive to the survival of parasitic nematodes. In contrast, classically activated macrophages appear to be deleterious to the survival of Taeniid larvae, occurring in the resistant Th1 acute phase of infection, whilst their phenotype is shifted to an alternatively activated state during chronic Taeniid infection (8,52). This could potentially explain why it may not be beneficial for Taeniids to secrete large amounts of AChE.

Nonetheless, our findings demonstrate that both *T. solium* and *T. crassiceps* larvae do contain AChEs, and using whole-cell patch clamp recordings in rodent hippocampal brain slice cultures, we show that *Taenia* larval-derived AChEs have sufficient activity to modify neuronal responses to acetylcholine. This has implications in the context of neurocysticercosis. Neurocysticercosis often presents with an extended asymptomatic period while the cyst remains viable, and it is only when the cyst loses viability and subsequently begins to disintegrate that symptom onset commonly occurs (53). It seems probable that during this phase of cyst degeneration, components of the cyst vesicular fluid, cyst membrane, and scolex that would otherwise be separated from the brain tissue by the tegumental membrane come into contact with the brain parenchymal cells. The exposure of brain tissue to larval AChEs during cyst degradation could interfere with endogenous cholinergic signalling. Here we have shown that AChEs reduced acetylcholine-induced pyramidal cell depolarization, which is an inhibitory action. Interestingly however, Zimmerman *et al.* (2008) have reported that in epileptic rats, there is a shift from membrane bound AChEs to soluble, unbound AChEs, and that this is associated with an increased sensitivity to acetylcholine, which in turn results in acetylcholine signalling inducing seizure activity in the epileptic animals. It seems conceivable, then, that an increase in soluble AChEs in the extracellular brain environment as a result of cyst degeneration could similarly induce greater acetylcholine sensitivity and could contribute to the generation of seizure activity in neurocysticercosis, particularly if combined with other seizure-promoting processes.

Additional epileptogenic processes in neurocysticercosis very likely involve the neuroinflammatory response, as there is mounting evidence that brain inflammatory processes contribute to the generation and subsequent development of seizures and epilepsy (55,56). Furthermore, the degeneration of *T. solium* cysts is associated with a strong pro-inflammatory host immune response, which is typically correlated with seizure onset or an aggravation of symptoms (57,58). Microglia and astrocytes regulate inflammatory signalling in the brain via, amongst others, acetylcholine receptor dependent signalling (59). Activation of acetylcholine receptors can strikingly impair acute phase inflammation (60,61). This supports the presence of parasitic AChEs driving exacerbated inflammation around the lesion by reducing acetylcholine receptor mediated antiinflammatory signalling. This could further contribute to perilesional gliosis which occurs in a subset of neurocysticercosis cases, and is associated with seizure recurrence (62). Future work, possibly utilising *in vivo* animal models, would be necessary to provide more definitive evidence implicating larval derived AChEs in neurocysticercosis-associated epilepsy.

In summary, our findings describe distinct profiles of acetylcholinesterase activity in *T. crassiceps* and *T. solium* larvae. In so doing, we highlight the possibility of larval-derived enzymes interfering with both host neural and immune signalling in the brain.

## Acknowledgements

*T. crassiceps* larvae were generously donated to us by Dr Siddhartha Mahanty (University of Melbourne, Melbourne, Australia).

## References

1. Garcia HH, Gonzalez AE, Gilman RH. Taenia solium Cysticercosis and Its Impact in Neurological Disease. Clin Microbiol Rev. 2020;33(3):1–23.

2. Carpio A, Romo ML. Multifactorial basis of epilepsy in patients with neurocysticercosis. Epilepsia. 2015;56(6):973–82.

3. Nash TE, Mahanty S, Garcia HH. Neurocysticercosis-More Than a Neglected Disease. PLoS Negl Trop Dis. 2013;7(4):7–9.

4. Roman G, Sotelo J, Del Brutto O, Flisser A, Dumas M, Wadia N, et al. A proposal to declare neurocysticercosis an international reportable disease. Bull World Health Organ. 2000;78(3):399–406.

5. Bhattarai R, Budke CM, Carabin H, Proaño J V., Flores-Rivera J, Corona T, et al. Quality of life in patients with neurocysticercosis in Mexico. Am J Trop Med Hyg. 2011;84(5):782–6.

6. Dzik JM. Molecules released by helminth parasites involved in host colonization. Acta Biochim Pol. 2006;53(1):33–64.

7. Mcsorley HJ, Maizels RM. Helminth Infections and Host Immune Regulation. Clin Microbiol Rev. 2012;25(4):585–608.

8. Peon AN, Ledesma-Soto Y, Terrazas LI, Peón AN, Ledesma-Soto Y, Terrazas LI, et al. Regulation of immunity by Taeniids: lessons from animal models and in vitro studies. Parasite Immunol. 2016;38(October 2015):124–35.

9. Leflore WB, Smith BF. The Histochemical Localization of Esterases in Whole Mounts of Cysticercus fasciolaris. Trans Am Microsc Soc. 1976;95(1):73–9.

10. Selkirk ME, Lazari O, Matthews JB. Functional genomics of nematode acetylcholinesterases. Parasitology. 2005;131:S3–18.

11. Schwabe CW, Koussa M, Acra AN. Host-parasite relationships in echinococcosis – IV. Acetylcholinesterase and permeability regulation in the Hydatid cyst wall. Comp Biochem Physiol. 1961;2:161–72.

12. Espinoza B, Tarrab-Hazdai R, Himmeloch S, Arnon R. Acetylcholinesterase from Schistosoma mansoni: immunological characterization. Immunol Lett. 1991;28(2):167–74.

13. Gimenez-Pardo C, Martinez-Grueiro MM, Gomez-Barrio A, Martinez-Fernandez AR, Rodriguez-Caabeiro F. Phosphomonoesterases and cholinesterases from Taenia pisiformis cysticerci. Helminthologia. 2004;3:115–20.

14. Camacho M, Tarrab-Hazdai R, Espinoza B, Arnon R, Agnew A. The amount of acetylcholinesterase on the parasite surface reflects the differential sensitivity of schistosome species to metrifonate. Parisitology. 1994;108:153–60.

15. Vaux R, Schnoeller C, Berkachy R, Roberts LB, Hagen J, Gounaris K, et al. Modulation of the Immune Response by Nematode Secreted Acetylcholinesterase Revealed by Heterologous Expression in Trypanosoma musculi. PLoS Pathog. 2016;12(11: e1005998.):1–18.

16. Tedla BA, Sotillo J, Pickering D, Eichenberger RM, Ryan S, Becker L, et al. Novel cholinesterase paralogs of Schistosoma mansoni have perceived roles in cholinergic signaling and drug detoxification and are essential for parasite survival [Internet]. Vol. 15, PLoS Pathogens. 2019. 1–32 p. Available from: http://dx.doi.org/10.1371/journal.ppat.1008213

17. Darby M, Schnoeller C, Vira A, Culley F, Bobat S, Logan E, et al. The M3 Muscarinic Receptor Is Required for Optimal Adaptive Immunity to Helminth and Bacterial Infection. PLoS Pathog. 2015;11(1):1–15.

18. Friedman A, Behrens CJ, Heinemann U. Pathophysiology of Chronic Epilepsy: Cholinergic Dysfunction in Temporal Lobe Epilepsy. Epilepsia. 2007;48(Suppl.:126–30.

19. Colangelo C, Shichkova P, Keller D, Markram H, Ramaswamy S. Cellular, Synaptic and Network Effects of Acetylcholine in the Neocortex. Front Neural Circuits. 2019;13(April):Article 24.

20. Gotti C, Zoli M, Clementi F. Brain nicotinic acetylcholine receptors: native subtypes and their relevance. Trends Pharmacol Sci. 2006;27(9):482–91.

21. Picciotto MR, Higley MJ, Mineur YS. Acetylcholine as a Neuromodulator: Cholinergic Signaling Shapes Nervous System Function and Behavior. Neuron [Internet]. 2012;76(1):116–29. Available from: http://dx.doi.org/10.1016/j.neuron.2012.08.036

22. Raggenbass M, Bertrand D. Nicotinic Receptors in Circuit Excitability and Epilepsy. Wiley Intersci. 2002;(May):580–9.

23. Curia G, Longo D, Biagini G, Jones RSG, Avoli M. The pilocarpine model of temporal lobe epilepsy. J Neurosci Methods. 2008;172:143–57.

24. Cordner SM, Fysh RR, Gordon H. Deaths of two hospital inpatients poisoned by pilocarpine. Br Med J. 1986;293(November):1285–7.

25. Tattersall J. Seizure activity post organophosphate exposure. Front Biosci. 2009;14:3688–711.

26. Ntoukas V, Tappe D, Pfütze D, Simon M, Holzmann T. Cerebellar Cysticercosis Caused by Larval Taenia crassiceps Tapeworm in Immunocompetent Woman, Germany. Emerg Infect Dis. 2013;19(12):2008–11.

27. de Lange A, Mahanty S, Raimondo J V. Model systems for investigating disease processes in neurocysticercosis. Parasitology [Internet]. 2018;146(5):553–62. Available from: https://www.cambridge.org/core/product/identifier/S0031182018001932/type/journal_article

28. Shield JM. Dipylidium caninum, Echinococcus granulosus and Hydatigera taeniaeformis: Histochemical Identification of Cholinesterases. Exp Parasitol. 1969;231:217–31.

29. Eranko O, Kouvalainen K, Mattila M, Takki S. Histochemical and Biochemical Observations on Cholinesterases of Cat’s Tapeworm Taenia Taeniaformis. Acta Physiol Scand. 1968;73:226–33.

30. Lee DL, Rothman AH, Senturia JB. Esterases in Hymenolepis and in Hydatigera. Exp Parasitol. 1963;14:285–95.

31. Cumino AC, Nicolao MC, Loos JA, Denegri G, Elissondo MC. Echinococcus granulosus tegumental enzymes as in vitro markers of pharmacological damage: A biochemical and molecular approach. Parasitol Int [Internet]. 2012;61(4):579–85. Available from: http://dx.doi.org/10.1016/j.parint.2012.05.007

32. Gimenez-Pardo C, Ros Moreno RM, de Armas-Serra C, Rodriguez-Caabeiro F. Presence of cholinesterase in Exhinococcus Granulosis protoscolices. Parasite. 2000;7:47–50.

33. Ciurea A V, Fountas KN, Coman TC, Machinis TG, Kapsalaki EZ, Fezoulidis NI, et al. Long-term surgical outcome in patients with intracranial hydatid cyst. Acta Neurochir (Wien). 2006;148:421–6.

34. Koziol U, Krohne G, Brehm K. Anatomy and development of the larval nervous system in Echinococcus multilocularis. Front Zool. 2013;10(1):1–17.

35. Trejo-Chávez H, García-Vilchis D, Reynoso-Ducoing O, Ambrosio JR. In vitro evaluation of the effects of cysticidal drugs in the Taenia crassiceps cysticerci ORF strain using the fluorescent CellTracker CMFDA. Exp Parasitol. 2011;127:294–9.

36. Vasantha S, Ravi Kumar B V, Roopashree SD, Das S, Shankar SK. Neuroanatomy of Cysticercus cellulosae (Cestoda) as revealed by acetylcholinesterase and nonspecific esterase histochemistry. Parasitol Res. 1992;78:581–6.

37. Martínez-Zedillo G, González-Barranco, D González-Angulo A. Presence of esterases and peptidases in the intact tegument of vesicles of Cysticercus cellulosae. Arch Invest Med. 1983;14:367–77.

38. Parija SC, Ar G. Cysticercus cellulosae antigens in the serodiagnosis of neurocysticercosis. Trop Parasitol. 2011;1(2):64–72.

39. Tomes H, de Lange A, Prodjinotho UF, Mahanty S, Smith K, Horsnell W, et al. Cestode larvae excite host neuronal circuits via glutamatergic signaling – preprint. bioRxiv. 2020;

40. Ellman GL, Courtney KD, Andres V, Featherstone RM. A new and rapid colorimetric determination of acetylcholinesterase activity. Biochem Pharmacol. 1961;7:88–95.

41. Selkirk ME, Hussein AS. Acetylcholinesterases of gastrointestinal nematodes. In: Chudi C, Pearce EJ, editors. Biology of parasitism: A modern approach. Kluwer Academic Publishers; 2000. p. 121–42.

42. Karnovsky MJ, Roots L. A “direct-coloring” thiocholine method for cholinesterases. J Histochem Cytochem. 1964;12:219–21.

43. Stoppini L, Buchs P-A, Muller D. A simple method for organotypic cultures of nervous tissue. J Neurosci Methods. 1991;37:173–82.

44. Forman CJ, Tomes H, Mbobo B, Burman RJ, Jacobs M, Baden T, et al. Openspritzer: An open hardware pressure ejection system for reliably delivering picolitre volumes. Sci Rep. 2017;7(2188).

45. Austin L, Berry WK. Two selective inhibitors of cholinesterase. Biochem J. 1953;54(4):695–700.

46. Cole BYAE, Nicoll RA. Characterization of a slow cholinergic post-synaptic potential recorded in vitro from rat hippocampal pyramidal cells. J Physiol. 1984;352:173–88.

47. Robinson P, Garza A, Weinstock J, Serpa JA, Goodman JC, Eckols KT, et al. Substance P Causes Seizures in Neurocysticercosis. PLoS Pathog. 2012;8(2):e1002489.

48. Stringer JL, Marks LM, White, Jr. AC, Robinson P. Epileptogenic activity of granulomas associated with murine cysticercosis. Exp Neurol. 2003;183:532–6.

49. Matos-Silva H, Reciputti BP, De Paula ÉC, Oliveira AL, Moura VBL, Vinaud MC, et al. Experimental encephalitis caused by Taenia crassiceps cysticerci in mice. Arq Neuropsiquiatr. 2012;70(4):287–92.

50. Leandro LDA, Fraga CM, de Souza Lino Jr R, Vinaud MC. Partial reverse of the TCA cycle is enhanced in Taenia crassiceps experimental neurocysticercosis after in vivo treatment with anthelminthic drugs. Parasitol Res. 2014;113:1313–7.

51. Kemmerling U, Cabrera G, Campos EO, Inestrosa NC, Galanti N. Localization, specific activity, and molecular forms of acetylcholinesterase in developmental stages of the cestode Mesocestoides corti. J Cell Physiol. 2006;206(2):503–9.

52. Prodjinotho UF, Lema J, Lacorcia M, Schmidt V, Vejzagic N, Sikasunge C, et al. Host immune responses during Taenia solium Neurocysticercosis infection and treatment. PLoS Negl Trop Dis [Internet]. 2020;14(4):e0008005. Available from: http://dx.doi.org/10.1371/journal.pntd.0008005

53. White, Jr. AC. Neurocysticercosis: Updates on Epidemiology, Pathogenesis, Diagnosis, and Management. Annu Rev Med. 2000;51:187–206.

54. Zimmerman G, Njunting M, Ivens S, Tolner E, Behrens CJ, Gross M, et al. Acetylcholine-induced seizure-like activity and modified cholinergic gene expression in chronically epileptic rats. Eur J Neurosci. 2008;27(4):965–75.

55. Vezzani A. Epilepsy and Inflammation in the Brain: Overview and Pathophysiology. Epilepsy Curr. 2014;14(1):3–7.

56. de Vries EE, van den Munckhof B, Braun KPJ, van Royen-Kerkhof A, de Jager W, Jansen FE. Inflammatory mediators in human epilepsy: A systematic review and meta-analysis. Neurosci Biobehav Rev [Internet]. 2016;63:177–90. Available from: http://dx.doi.org/10.1016/j.neubiorev.2016.02.007

57. Uddin J, Gonzalez AE, Gilman RH, Thomas LH, Rodriguez S, Evans CAW, et al. Mechanisms Regulating Monocyte CXCL8 Secretion in Neurocysticercosis and the Effect of Antiparasitic Therapy. J Immunol. 2010;185(7):4478–84.

58. Singh AK, Prasad KN, Prasad A, Tripathi M, Gupta RK, Husain N. Immune responses to viable and degenerative metacestodes of Taenia solium in naturally infected swine. Int J Parasitol [Internet]. 2013;43(14):1101–7. Available from: http://dx.doi.org/10.1016/j.ijpara.2013.07.009

59. Carnevale D, De Simone R, Minghetti L. Microglia-Neuron Interaction in Inflammatory and Diseases: Role of Cholinergic and Noradrenergic Systems. CNS Neurol Disord – Drug Targets. 2007;6:388–97.

60. Patel H, Mcintire J, Ryan S, Dunah A, Loring R. Anti-inflammatory effects of astroglial α7 nicotinic acetylcholine receptors are mediated by inhibition of the NF-κB pathway and activation of the Nrf2 pathway. J Neuroinflammation. 2017;14(192):1–15.

61. Binning W, Hogan-Cann AE, Sakae DY, Maksoud M, Ostapchenko V, Al-Onaizi M, et al. Chronic hM3Dq signaling in microglia ameliorates neuroinflammation in male mice – preprint. bioRxiv. 2020;

62. Pradhan S, Kathuria MK, Gupta RK. Perilesional Gliosis and Seizure Outcome: A Study Based on Magnetization Transfer Magnetic Resonance Imaging in Patients with Neurocysticercosis. Ann Neurol. 2000;48(2):181–7.

