## Supplemental tables for "*Taenia* larvae possess distinct acetylcholinesterase profiles with implications for host cholinergic signalling"

**Supplementary Table 1:** Sensitivity of *T. crassiceps* cholinesterase activity in various extracts to different inhibitors.

| | | | Inhibitor concentration | | | | | |
|---|---|---|---|---|---|---|---|---|
| | | | **Uninhibited** | **100 nM** | **1 µM** | **10 µM** | **100 µM** | **1 mM** |
| | | | **Cholinesterase activity (% of mean uninhibited)** | | | | | |
| **BW284C51** | Whole cyst homogenate | Median | 100.4 | 99.4 | 98.9 | 84.8 | 21.3 | 2.2 |
| | | Range | 96.0 – 104.0 | 95.1 – 108.3 | 98.8 – 101.7 | 83.9 – 85.6 | 18.8 – 22.8 | 1.3 – 4.1 |
| | | # of assays | 10 | 3 | 3 | 3 | 3 | 3 |
| | Cyst membrane | Median | 100.5 | 94.0 | 96.8 | 75.3 | 28.7 | 3.6 |
| | | Range | 81.6 – 109.2 | 93.2 – 95.3 | 73.4 – 97.0 | 64.5 – 78.9 | 28.4 – 30.1 | 2.9 – 4.3 |
| | | # of assays | 10 | 3 | 3 | 3 | 3 | 3 |
| | Cyst vesicular fluid | Median | 100.5 | 96.3 | 93.2 | 72.6 | 25.5 | 5.2 |
| | | Range | 97.0 – 101.4 | 94.3 – 96.8 | 92.4 – 93.4 | 72.5 – 79.2 | 25.2 – 30.8 | 5.1 – 5.3 |
| | | # of assays | 10 | 3 | 3 | 3 | 3 | 3 |
| | Excretory/ secretory products | Median | 99.7 | 85.9 | 93.3 | 73.2 | 21.5 | 2.2 |
| | | Range | 97.6 – 105.1 | 85.6 – 90.1 | 93.1 – 98.3 | 70.6 – 75.7 | 19.5 – 21.8 | UD – 4.4 |
| | | # of assays | 10 | 3 | 3 | 3 | 3 | 3 |
| **Iso-OMPA** | Whole cyst homogenate | Median | 100.4 | 96.2 | 97.5 | 98.4 | 105.0 | 82.3 |
| | | Range | 96.0 – 104.0 | 91.7 – 100.4 | 91.2 – 101.0 | 96.6 – 101.2 | 94.1 – 113.9 | 19.1 – 100.8 |
| | | # of assays | 10 | 3 | 5 | 3 | 5 | 5 |
| | Cyst membrane | Median | 100.5 | 94.6 | 101.9 | 97.8 | 96.2 | 83.2 |
| | | Range | 81.6 – 109.2 | 94.1 – 98.0 | 92.5 – 107.2 | 97.1 – 100.2 | 75.0 – 116.9 | 80.2 – 91.5 |
| | | # of assays | 10 | 3 | 5 | 3 | 5 | 5 |
| | Cyst vesicular fluid | Median | 100.5 | 97.0 | 96.1 | 98.9 | 99.3 | 96.3 |
| | | Range | 97.0 – 101.4 | 96.4 – 97.4 | 95.0 – 97.5 | 98.0 – 99.1 | 97.3 – 101.3 | 89.4 – 103.0 |
| | | # of assays | 10 | 3 | 5 | 3 | 5 | 5 |
| | Excretory/ secretory products | Median | 99.7 | 103.7 | 99.9 | 103.1 | 100.6 | 96.8 |
| | | Range | 97.5 – 105.1 | 102.9 – 105.7 | 96.7 – 104.3 | 99.5 – 106.0 | 95.0 – 111.2 | 87.3 – 106.6 |
| | | # of assays | 10 | 3 | 5 | 3 | 5 | 5 |
| **Eserine** | Whole cyst homogenate | Median | 100.4 | 52.5 | 55.4 | 49.6 | UD | UD |
| | | Range | 96.0 – 104.0 | 52.1 – 54.4 | 53.9 – 56.4 | 49.1 – 50.2 | UD | UD |
| | | # of assays | 10 | 3 | 3 | 3 | 3 | 3 |
| | Cyst membrane | Median | 100.5 | 52.1 | 5.8 | 1.8 | 2.1 | UD |
| | | Range | 81.6 – 109.2 | 49.3 – 53.7 | 4.7 – 6.2 | 0.8 – 2.1 | 1.8 – 3.2 | UD |
| | | # of assays | 10 | 3 | 3 | 3 | 3 | 3 |
| | Cyst vesicular fluid | Median | 100.5 | 56.4 | 8.1 | 3.8 | 4.1 | 0.7 |
| | | Range | 97.0 – 101.4 | 54.8 – 56.4 | 6.8 – 9.3 | 2.4 – 4.9 | 3.8 – 4.6 | 0.5 – 1.3 |
| | | # of assays | 10 | 3 | 3 | 3 | 3 | 3 |
| | Excretory/ secretory products | Median | 99.7 | 47.3 | 8.5 | 2.8 | UD | 0.4 |
| | | Range | 97.5 – 105.1 | 46.3 – 49.5 | 5.3 – 8.8 | 0.7 – 4.2 | UD | 0.3 – 2.1 |
| | | # of assays | 10 | 3 | 3 | 3 | 3 | 3 |

UD = undetectable, where activity was so low as to be undetectable.

**Supplementary Table 2:** Sensitivity of *T. solium* cholinesterase activity in various extracts to different inhibitors.

| | | | Inhibitor concentration | | | | | |
|---|---|---|---|---|---|---|---|---|
| | | | **Uninhibited** | **100 nM** | **1 µM** | **10 µM** | **100 µM** | **1 mM** |
| | | | **Cholinesterase activity (% of mean uninhibited)** | | | | | |
| **BW284C51** | Whole cyst homogenate | Median | 105.3 | - | 99.5 | 58.7 | 15.3 | 9.9 |
| | | Range | 73.7 – 110.5 | - | 97.8 – 102.0 | 55.3 – 66.3 | 12.8 – 17.9 | 8.2 – 9.9 |
| | | # of assays | 5 | - | 3 | 5 | 3 | 3 |
| | Cyst membrane & scolex | Median | 96.9 | - | 136.8 | 57.9 | 0.0 | - |
| | | Range | 95.2 – 107.1 | - | 131.6 – 142.1 | 52.6 – 97.4 | 0.0 – 7.9 | - |
| | | # of assays | 5 | - | 3 | 5 | 3 | - |
| | Cyst vesicular fluid | Median | 108.6 | - | 118.6 | 42.9 | 0.0 | - |
| | | Range | 54.3 – 122.9 | - | 104.3 – 121.4 | 21.4 – 50.0 | 0.0 – 2.9 | - |
| | | # of assays | 5 | - | 3 | 5 | 3 | - |
| **Iso-OMPA** | Whole cyst homogenate | Median | 105.3 | - | 106.3 | 107.1 | 86.7 | - |
| | | Range | 73.7 – 110.5 | - | 103.7 – 112.2 | 100.3 – 113.1 | 85.8 – 92.7 | - |
| | | # of assays | 5 | - | 3 | 5 | 3 | - |
| | Cyst membrane & scolex | Median | 96.9 | - | 128.9 | 118.4 | 84.2 | - |
| | | Range | 95.2 – 107.1 | - | 107.9 – 152.6 | 107.9 – 152.6 | 84.2 – 86.8 | - |
| | | # of assays | 5 | - | 3 | 5 | 3 | - |
| | Cyst vesicular fluid | Median | 108.6 | - | 108.6 | 95.7 | 88.6 | - |
| | | Range | 54.3 – 122.9 | - | 102.9 – 122.9 | 85.7 – 112.9 | 82.9 – 90.0 | - |
| | | # of assays | 5 | - | 3 | 5 | 3 | - |
| **Eserine** | Whole cyst homogenate | Median | 105.3 | 96.9 | 49.3 | 47.6 | 19.6 | UD |
| | | Range | 73.7 – 110.5 | 95.2 – 107.1 | 49.3 – 51.0 | 46.8 – 50.2 | 9.4 – 28.1 | UD |
| | | # of assays | 5 | 5 | 3 | 3 | 5 | 3 |
| | Cyst membrane & scolex | Median | 96.9 | 105.3 | 47.4 | 97.4 | 44.7 | UD |
| | | Range | 95.2 – 107.1 | 73.7 – 110.5 | 44.7 – 47.4 | 60.0 – 102.6 | 7.9 – 63.2 | UD |
| | | # of assays | 5 | 5 | 3 | 3 | 5 | 3 |
| | Cyst vesicular fluid | Median | 108.6 | 20.0 | 72.8 | 50.0 | UD | - |
| | | Range | 54.3 – 122.9 | 11.4 – 34.3 | 67.1 – 110.0 | 38.6 – 64.3 | UD | - |
| | | # of assays | 5 | 5 | 3 | 3 | 5 | - |

UD = undetectable, where activity was so low as to be undetectable.
